# A natural language fMRI dataset for voxelwise encoding models

**DOI:** 10.1101/2022.09.22.509104

**Authors:** Amanda LeBel, Lauren Wagner, Shailee Jain, Aneesh Adhikari-Desai, Bhavin Gupta, Allyson Morgenthal, Jerry Tang, Lixiang Xu, Alexander G. Huth

## Abstract

Speech comprehension is a complex process that draws on humans’ abilities to extract lexical information, parse syntax, and form semantic understanding. These sub-processes have traditionally been studied using separate neuroimaging experiments that attempt to isolate specific effects of interest. More recently it has become possible to study all stages of language comprehension in a single neuroimaging experiment using narrative natural language stimuli. The resulting data are richly varied at every level, enabling analyses that can probe everything from spectral representations to high-level representations of semantic meaning. We provide a dataset containing BOLD fMRI responses recorded while 8 subjects each listened to 27 complete, natural, narrative stories (~6 hours). This dataset includes pre-processed and raw MRIs, as well as hand-constructed 3D cortical surfaces for each participant. To address the challenges of analyzing naturalistic data, this dataset is accompanied by a python library containing basic code for creating voxelwise encoding models. Altogether, this dataset provides a large and novel resource for understanding speech and language processing in the human brain.

## Background and Summary (Unlimited length)

Historically, MRI has been used to study the structural and functional organization of the brain by way of highly controlled paradigms and simplified stimuli. This has also been true in language neuroscience (i.e. using block designs^1,2^ where it is common to use isolated words^1,3^ or simple sentences^4–7^ as experimental stimuli. While these paradigms have proven useful, they also present several issues. First, their hypothesis-driven design limits the number of scientific questions one can ask using a given dataset. Second, isolated words and sentences are devoid of context, which is a critical component of language in the real world. Thus, results obtained on these simple stimuli may not generalize to natural language perception. And third, small stimulus sets limit the breadth of features sampled by the stimuli. This is problematic for studying complex, high-dimensional feature spaces such as semantics.

An alternative approach is to use natural stimuli that closely approximate language as it is used in everyday life. *Natural language* is language used in real world settings such as conversation, entertainment, and education. Natural language can also include multiple modalities, such as vision for written or signed language and audition for spoken language.^8^ Our dataset focuses on one specific subset of natural language: spoken English in the form of complete narrative stories from *The Moth* podcast. This permits detailed study of the auditory speech processing as well as core amodal language systems. While narrow compared to the full breadth of natural language, these stories are highly varied in semantic and syntactic content. There is also considerable evidence from earlier experiments that such natural stories broadly activate cortex and can be used to study a variety of phenomena.^9–16^

One downside of natural language stimuli is the difficulty of analyzing and interpreting the resulting data. Data from controlled, block-design experiments can be analyzed using standard methods such as t-tests, f-tests, and ANOVAs. In natural language, however, the features of interest (e.g. a particular phoneme, or topic) are distributed throughout the stimulus. To model how the brain responds to these features—and account for correlations among them—we use *voxelwise encoding models* that are designed to predict brain responses from the stimuli^17^. The first step in creating encoding models is to extract features of interest from the stimuli. Previous work on this fMRI dataset has used spectral, articulatory, part-of-speech, and semantic features to predict BOLD responses.^9,18–20^ After features are extracted for each word, phoneme, or timepoint, they are downsampled to the rate of the fMRI acquisition, delayed to account for hemodynamic response, and used in a regression model to predict the fMRI data. These models are fit separately for each voxel in each subject, providing high resolution and high fidelity.

For this dataset, we conducted an fMRI experiment in which eight participants passively listened to 27 complete, natural, narrative stories (370 minutes) from *The Moth Radio Hour* over the course of five scanning sessions. While only covering a few participants, the large amount of data per participant enables more sophisticated analyses than would be possible with fewer stimuli. Each story was transcribed, aligned, and hand-checked to provide the timing of every word and phoneme. Functional localizer data for known sensorimotor, auditory, and cognitive regions was also collected, as well as high resolution T1-weighted structural scans which were used to create hand-corrected cortical surfaces for each participant. For a summary of available resources see **Figure 1**.

**Figure 1.**
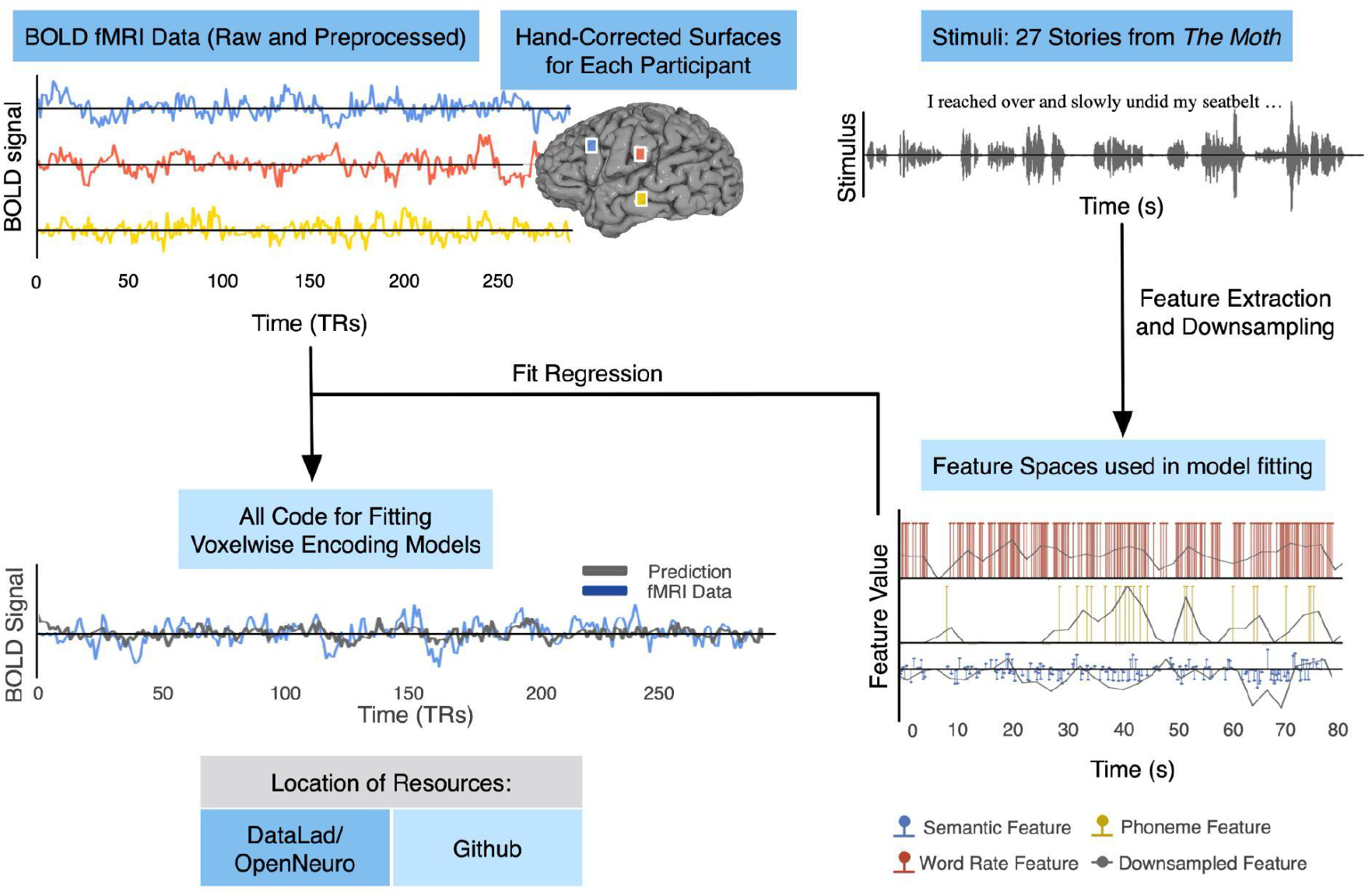
*Schematic of naturalistic story-listening paradigm and available resources.* 27 unique natural stories from *The Moth* podcast were played for eight participants over five fMRI sessions while they were instructed to passively listen. One of these 27 stories was played in each of the 5 sessions. No other story was repeated. These stimuli can be converted to previously used feature spaces for model fitting, including semantic, phoneme, and word rate feature spaces.^9,10,13^ Regularized regression can then be used to fit voxelwise encoding models that use the features to predict BOLD data. Model performance can then be evaluated on a held out dataset. Available resources on OpenNeuro include the stimuli, BOLD data, and hand-corrected surfaces for each of the eight participants. Available resources on GitHub include the feature spaces and code for fitting the encoding models.

Voxelwise models are typically evaluated on a separate test dataset. Trained encoding models and test stimuli are used to predict response time courses for each voxel. Prediction performance is then computed as the correlation between predicted and actual responses. This type of evaluation, which tests how well a model can predict responses to novel natural stimuli, is a good proxy for how well the model captures language representations in the brain.^21^ Here, the test dataset comprises one story, which was played once in each of the five scanning sessions. These five responses can be averaged to obtain a high-quality test dataset, or they can be used to de-bias the prediction performance measure through noise-ceiling estimation.^22^

## Methods (unlimited length)

### Participants

Data was collected from 8 participants (three female): UTS01 (female, age 24), UTS02 (male, age 34), UTS03 (male, age 21), UTS04 (male, age 31), UTS05 (female, age 24), UTS06 (female, age 23), UTS07 (male, age 25), UTS08 (male, age 24). All participants were healthy and had normal hearing. The experimental protocol was approved by the Institutional Review Board at the University of Texas at Austin, and written informed consent was obtained from all participants.

### Natural Language Stimulus Set

The stimulus set consisted of 26 10-15 minute stories (320 minutes and 24 seconds total duration) from *The Moth Radio Hour* plus one additional 10-minute story played in each session to be used as a test dataset (50 minutes), giving a total of 370 minutes of data per subject. For stimulus presentation, the audio for each story was filtered to correct for frequency response and phase errors induced by the headphones using calibration data provided by Sensimetrics and custom Python code (https://github.com/alexhuth/sensimetrics_filter). All stimuli were played at 44.1 kHz using the pygame library in Python.

In each story, a single speaker tells an autobiographical story without reading from a prepared script. All stories were manually transcribed by one listener. The Penn Phonetics Lab Forced Aligner (P2FA)^27^ was then used to automatically align the audio to the transcript. Certain sounds (for example, laughter and breathing) were also marked to improve the accuracy of the automated alignment (see **Table 1**). Praat^28^ phonetic analysis software was used to manually check and correct the alignment of each word within the transcript.

**Table 1:**
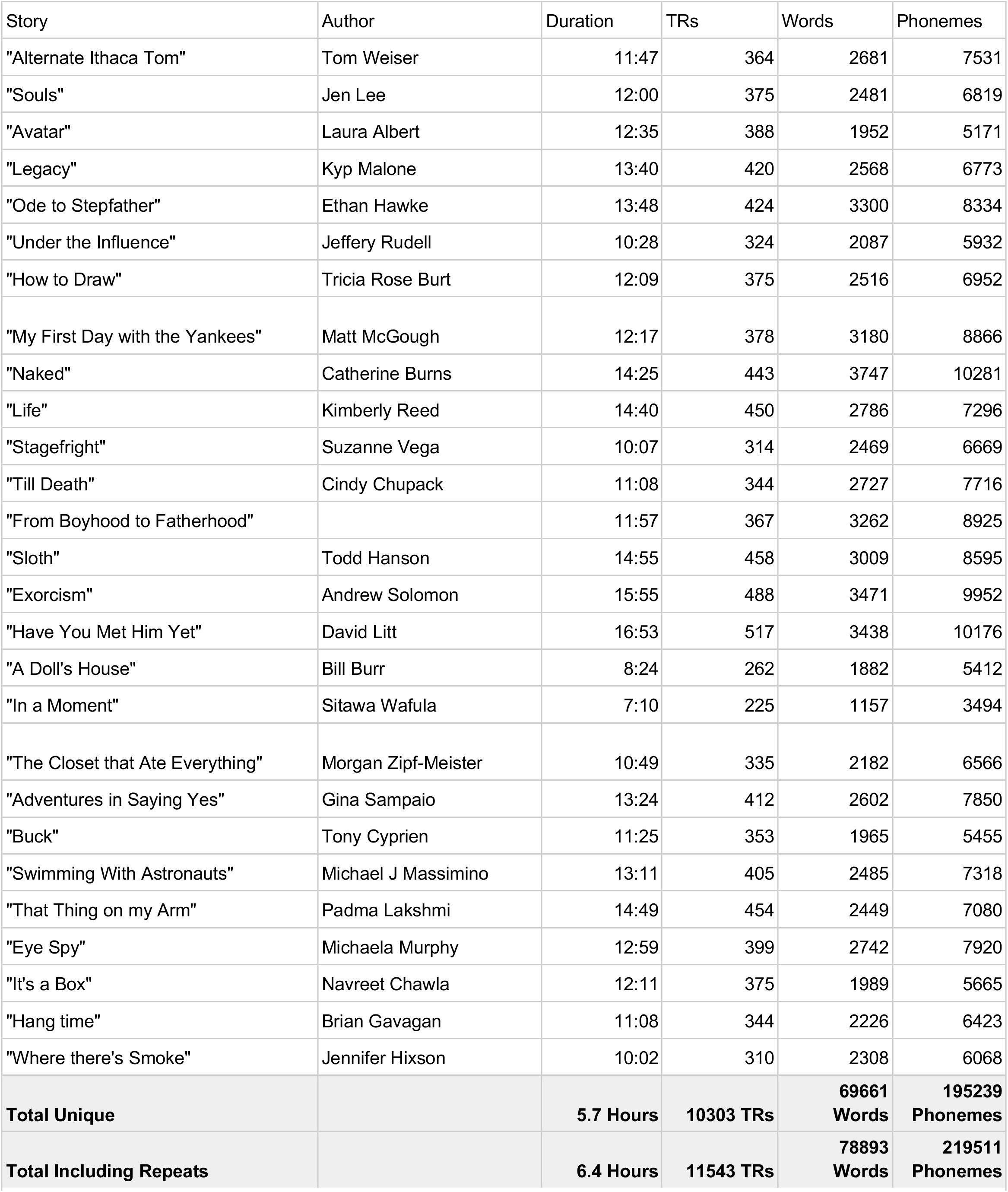
Story stimulus from the *Moth Radio Hour.* All story stimuli were from the Moth Radio Hour and were hand transcribed and the transcipts were then aligned to the audio. For each story, we have included the original author and speaker for each narrative story and the total length by time, TRs, words and phonemes in the stimuli.

**Table 2:**
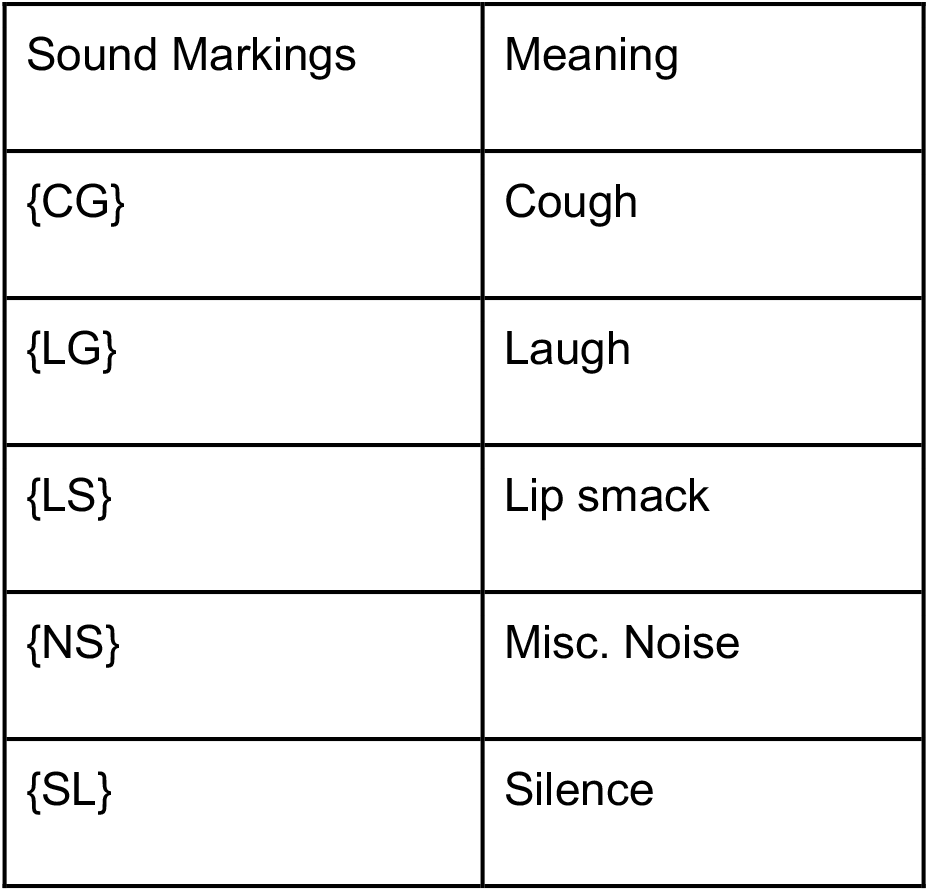
Additional sounds marked in scripts. Certain sounds were included in hand aligned transcripts of the stimulus. This was done to improve alignment from the transcripts to the audio.

| Sound Markings | Meaning |
| --- | --- |
| {CG} | Cough |
| {LG} | Laugh |
| {LS} | Lip smack |
| {NS} | Misc. Noise |
| {SL} | Silence |

### fMRI data collection

MRI data was collected over 6 scanning sessions on a 3T Siemens Skyra scanner at the UT Austin Biomedical Imaging Center using a 64-channel Siemens volume coil. Anatomical data for participant UT-S-02 was collected on a 3T Siemens TIM Trio at the Berkeley Brain Imaging Center using a 32-channel Siemens volume coil. The first session included an anatomical scan and functional localizers. Each subsequent session consisted of passively listening to 4-5 stories, plus the story used for model testing. Each story was played during one EPI scan that included padding of 10 seconds silence at the beginning and end of each story. Audio was delivered through Sensimetrics S14 in-ear piezoelectric headphones. To minimize head motion, foam headcases (CaseForge, Inc., now defunct) that precisely fill the space between the participant’s head and the headcoil were used during data collection. To create the headcases, an RGB Structure.io sensor (Occipital Inc.) was used to collect a 3-dimensional scan of each participant’s head while hair was compressed using a swim cap. These scans were then used to mill customized styrofoam headcases for each participant.

### fMRI parameters

Functional scans were collected using gradient-echo EPI with repetition time (TR) = 2.00 s, echo time (TE) = 30.8 ms, flip angle = 71°, multi-band factor (simultaneous multi-slice) = 2, voxel size = 2.6 mm x 2.6 mm x 2.6 mm (slice thickness = 2.6 mm), matrix size = (84, 84) and field of view = 220 mm. Field of view covered the entire cortex for all participants. Anatomical data was collected using a T1-weighted multi-echo MPRAGE sequence on the same 3T scanner with voxel size = 1 mm x 1 mm x 1 mm following the Freesurfer morphometry protocol.

### fMRI preprocessing

fMRI preprocessing was only done on the derivative data. This data was motion corrected using the FMRIB Linear Image Registration Tool (FLIRT) from FMRIB Software Library (FSL) 5.0^29^. After motion correction, all the volumes within each run were averaged to obtain a single template volume. Cross-run alignment was then performed by using FLIRT to align the template volume from each run to the template volume from the first run in the first story session. These automatic alignments were manually checked. The motion correction and cross-run transformations were then concatenated and used to resample the original data to a motion-corrected and cross-run-aligned space. This process avoids multiple resampling steps, thus minimizing unwanted blurring. Low frequency voxel response drift was then identified using a 2^nd^ order Savitzky-Golay filter with a 120-second window and subtracted from the signal. To avoid artifacts from onset transients and poor detrending performance at the edges of the data, responses were trimmed by removing 20 seconds (10 volumes) at the beginning and end of each scan. This removed the 10-second silent period as well as the first and last 10 seconds of each story. The mean response for each voxel was then subtracted and the remaining response scaled to have unit variance.

### Cortical surface reconstruction and visualization

All anatomical data has been defaced using pydeface. For the cortical surfaces, meshes were generated from the T1-weighted anatomical scans using FreeSurfer^30^. Before surface reconstruction, anatomical surface segmentations were hand-checked and corrected. Blender *(https://blender.org)* was used to remove the corpus callosum and make relaxation cuts for flattening via the interface provided by pycortex^31^.

Functional images were aligned to the cortical surface using boundary-based registration (BBR) implemented in FSL. These were checked for accuracy and adjustments were made to the registration parameters as necessary.

Cortical maps of selectivity or model performance were created by projecting the values for each voxel onto the cortical surface using the ‘nearest’ scheme in pycortex^31^. This projection finds the location of each pixel in the image in 3D space, and assigns the pixel the value associated with the voxel enclosing that location.

### Functional Localizers and Region of Interest definitions

Known regions of interest (ROIs) were defined separately in each participant using three localizer tasks: a visual category localizer, a motor localizer, and an auditory cortex localizer. For the visual category localizer, data was collected in six 4.5 minute scans consisting of 16 blocks of 16 seconds each. During each block, 20 images of places, faces, bodies, household objects, or spatially scrambled objects were displayed. In order to encourage focus, participants were asked to perform a 1-back task where they pressed a button if the same image appeared twice in a row. The corresponding cortical ROIs defined with this localizer were the fusiform face area (FFA)^32^, occipital face area (OFA)^32^, extrastriate body area (EBA)^33^, parahippocampal place area (PPA)^34^, retrosplenial cortex (RSC), and the occipital place area (OPA). These ROIs were hand-drawn based on t-value maps from a contrasts comparing responses to faces and objects (FFA, OFA), bodies and objects (EBA), and places and objects (PPA, OPA, RSC).

The motor localizer data was collected during two identical 10-minute scans. The participant was cued to perform six different motor tasks in a random order in 20-second blocks. The cues ‘hand’, ‘foot’, ‘mouth’, ‘speak’, and ‘rest’ were visually presented at the center of the screen, and the *saccade* cue was presented as a random array of dots. For the *hand* cue, participants were instructed to make small finger-drumming movements for the duration of the cue. For the *foot* cue, participants were instructed to make small foot and toe movements. For the *mouth* cue, participants were instructed to make small nonsense vocalizations (i.e., the syllable string *“balabalabala”*). For the *speak* cue, participants were instructed to self-generate a narrative without vocalization. For the *saccade* cue, participants were instructed to look around for the duration of the task. Simple categorical regression models were fit for each voxel using ordinary least squares (OLS). Beta maps for the *hand, foot,* and *mouth* conditions were used to define primary motor and somatosensory areas for the hands, feet, and mouth; supplemental motor areas for the hands and feet; secondary motor areas for the hands, feet, and mouth; and the ventral premotor hand area (PMvh). The beta map for the *saccade* condition was used to define the frontal eye field and intraparietal sulcus visual areas. The beta map for the *speak* condition was used to define Broca’s area and the superior premotor ventral (sPMv) speech area^35^.

Auditory cortex localizer data was collected in one 10-minute scan. The participants listened to 10 repeats of a 1-minute auditory stimulus containing 20 seconds of music (Arcade Fire), speech (Ira Glass, This American Life), and nature sounds (a babbling brook). To determine whether a voxel was responsive to auditory stimulus, the repeatability of the voxel response across the 10 repeats was calculated using an F*-*statistic. This map was used to define the auditory cortex (AC).

### Stimulus Embeddings

The goal of encoding models is to find stimulus features that predict variance in the brain activity. This technique was originally developed for electrophysiology experiments^36^, but has now been used for BOLD signals in fMRI. In this framework, participants are presented with a stimulus, in this case stories, while brain activity is recorded. A linear regression model is then fit between some feature space, which is extracted from the stimulus, and the brain activity. The feature space serves as a hypothesis for the kind of information each voxel of brain data is representing. It is important to note that this is not a winner-take-all model and that each voxel is likely representing multiple different components of information.^17^

In this release we include python code for three feature spaces. The first is a word-level semantic feature space called *English1000*. This feature space has previously been used to map semantic representations across the cerebral cortex and cerebellum.^9,10,13^ English1000 is a 985-dimensional word embedding feature space based on word co-occurrence in English text.^10^ The second is an phoneme feature space. This is a 1-hot space comprising 44-dimensions, one for each phoneme in American English as defined by the CMU Pronouncing Dictionary as well as a few non-speech sounds. The last feature space is a word rate feature space. This is a 1dimensional feature space that represents the number of words spoken during each period of time. To create a feature space matrix for the regression, each word (or phoneme) in the stimulus is assigned a vector from the feature space. For example, for the stimulus phrase “I reached over”, one would take the embedding vectors for “I”, “reached”, and “over” from English1000 and concatenate them into a 3 (words) by 985 (features) matrix.

### Interpolation of the feature matrix

One challenge in fitting encoding models is that speech and BOLD data are sampled at very different frequencies. For example, approximately six words are spoken every two seconds, but only one brain image is recorded in that interval. To solve this problem, the stimulus matrix needs to be resampled to the same sampling frequency as the BOLD data. The procedure we provide for downsampling features to the fMRI acquisition rate can be thought of as comprising three steps. First, the discrete features for each word (or phoneme) are transformed into a continuous-time representation N(t) where t ∈[0, T] and T indicates the length of the stimulus. This representation is zero at all timepoints except for the exact middle of each word (or phoneme), where it is equal to an infinitesimal-duration spike (δ-function) that is scaled by the feature value. Next, a low-pass antialiasing Lanczos filter is convolved with N(t) to get N_LP_(t). The cutoff frequency of this antialiasing filter is selected to match the Nyquist frequency of the fMRI data (half the acquisition rate, or 0.25 Hz). Other parameters of the filter, such as the number of lobes, can be controlled in software. Finally, N_LP_(t) is sampled at the fMRI acquisition times t_r_ where r ∈ [1, 2,…n_TR_] corresponds to the volume index in the fMRI acquisition. In practice, these three steps are accomplished simultaneously by way of a single matrix multiplication: the word-(or phoneme-) level stimulus matrix *S* (number of features by number of words/phonemes) is multiplied by a sparse “Lanczos” matrix *L* (number of words/phonemes by number of fMRI volumes). In essence, this assumes that the total brain response is the sum of responses to each word or phoneme. This approach has been widely used for language encoding models with natural stimuli.^9,10,13,19,37,9,10,13,19^

### HRF Estimation

The BOLD responses recorded by fMRI are thought to capture delayed and low-pass filtered representations of local neural activity^38^. While most fMRI analyses treat the hemodynamic response function (HRF) as a fixed linear filter,^39^ the current dataset contains enough data that separate HRFs can be estimated for each voxel. To efficiently estimate individual HRFs, we use a finite impulse response (FIR) model^26^ in which separate model weights are estimated for each feature at several different delays (e.g. 2, 4, 6, and 8 seconds after the stimulus). This is accomplished by concatenating multiple versions of the interpolated stimulus matrix that have been delayed by different amounts.

### Model fitting

To fit the linear regression models that predict the response in each voxel, previous work using this dataset has used regularized regression. Regularized regression makes assumptions about the size or covariance of the regression weights. These assumptions constrain and improve the resulting models.^40^ Previous work on this and other natural language datasets has used L2-regularized or ridge regression.^10,17,41,42^ Ridge regression is computationally efficient and yields high model performance. We provide code for fitting ridge regression models as a part of the voxelwise modeling process.

### Model validation

To measure the predictive performance of an encoding model, it is important to use a held-out test data set. To do this, one takes the dot product of the regression weight matrix, consisting of a two-dimensional matrix of voxels by features, and the feature matrix of a new story not used in training the model (features by time). This results in a voxelwise prediction of brain activity in a 2-D matrix of voxels by time. The time-course of predicted brain response for each voxel is then correlated across time with the real brain data to measure model goodness-of-fit. It is important to use a held-out dataset for model evaluation, as failure to do so will result in overfitting and an overestimation of model performance. We recommend using the data elicited by *task-wheretheressmoke,* as this story was played multiple times for each participant. One issue is that the correlation between predicted and actual responses will be biased downwards due to noise in the fMRI data. The degree of bias can also vary from voxel to voxel depending on factors like proximity to vasculature,^23,24^ cortical folding,^24^ or other factors. To correct this bias, it is common to collect responses to the same test stimulus multiple times and then average them, decreasing the amount of noise in the test data.^10,12,13,18,25,26^ Averaging responses across repetitions effectively increases the signal to noise ratio of the BOLD response, providing less biased estimates of model performance

Another issue that can complicate interpretation of model performance values is that BOLD responses recorded using fMRI are inherently noisy, and the amount of noise can differ across brain areas and between participants. The amount of noise affects the maximum model performance that can be attained, even by a theoretically “perfect” model. To account for this one can attempt to estimate the noise ceiling of the data, which is the highest performance that any model can attain. This is done by comparing responses across repeats of the validation story, *task-wheretheressmoke.* We include code for computing the noise ceiling correction using a regularized normalized correlation coefficient (CC_norm_)^22^. This is done by first calculating the absolute product-moment correlation, defined as:

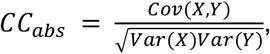

where X are the BOLD responses and Y are the model predictions. Then, to isolate model performance from prediction accuracy, this value is normalized as:

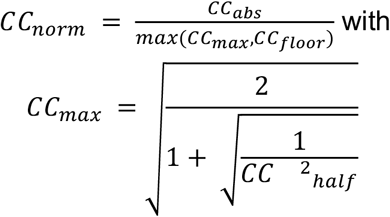

where CC_max_ is the noise ceiling. In comparison to the standard CC_norm_, we regularize the estimate by limiting the noise ceiling to be greater than CC_floor_=0.3. This value was determined to result in the least biased estimates in simulations using realistic noise values. Without this regularization, the estimated correlation after noise ceiling correction is not bounded and often surpasses a correlation of 1.0 for poorly-modeled voxels.

To test for significantly well-predicted voxels, one must compare model performance for each voxel to a null distribution. Here it is important to account for temporal auto-correlation in the BOLD signal. To do this, a block-wise permutation test is performed in which the true response timecourse is shuffled in blocks of 10 timepoints and then the correlation between model predictions and permuted data is computed. This process is repeated many times to form an empirical null distribution for each voxel. Block-wise permutation ensures that the permuted data retains temporal characteristics of the data while breaking the connection to the feature space being tested. Additionally, because each voxel is effectively treated as an independent model and each brain contains upwards of 80000 voxels, it is important to correct for multiple comparisons. In previous work, FDR correction was used to account for this^12,13,43^.

In prior studies, statistical significance was measured using prediction performance on one test story “task-wheretheressoke” that has multiple repeats.^9,10,12,13^ However, this approach also has a potential drawback: if a voxel is selective for features that are not present in the single test story, then those voxels may falsely be labeled as poorly predicted. To test for significance with multiple stories, an alternative approach is to use a leave-one-out procedure. This can be done by fitting an ensemble of encoding models, each of which excludes one unique training story (or session) in their model estimation. Statistical significance can then be measured for encoding model predictions on all held-out stories and their true BOLD responses, ie., the entire training set. This procedure increases diversity in the test set and improves statistical power.

## Data Records (unlimited length)

### Data

The raw data and derived data are available on OpenNeuro, including NIfTI files of all brain data, the story stimuli, derived data, hand-corrected surface reconstructions, and descriptions of paradigms. The data is organized into directories, one for each participant (8 total), chronologically organized by session. The first session includes both anatomical and functional data, broken up into corresponding folders. All other sessions only include functional data. Of note, the initial anatomical scan and functional localizer data for sub-UTS02 was collected previous to this dataset using different sequence parameters. Consequently, the localizer data is not included here. The BOLD data is stored in gzipped NIfTI 4D files under the naming pattern *sub-AA_ses-BB_task-CC_bold.nii.gz*. The story entitled *wheretheressmoke* is repeated in all five of the story sessions and thus also contains the run number in the name (i.e. *sub-AA_ses-BB_task-CC_run-DD_bold.nii.gz*). The preprocessed BOLD data is contained in HDF5 files for each story, organized by participant. The audio files for all of the story stimuli are stored as WAV files sampled at 44.1 kHz. Hand-corrected TextGrid files contain complete transcripts of each story as well as the temporal boundaries of each word and phoneme, and are stored as a derivative under Textgrids.

### Code

All of the standard code used to fit voxelwise encoding models is found at https://github.com/HuthLab/deep-fMRI-dataset.

‘encoding.py’ is the main script to train and evaluate encoding models. It takes X args (add mini explanation of each). First, the script finds the train and test stories for the specified fMRI session. Next, it loads the corresponding fMRI responses for the subject. It then obtains down-sampled stimulus features from ‘feature_spaces.py’ for every story, z-scores them and applies the FIR delays. It then trains and cross-validates a linear regression model on all the specified training stories. The returned encoding model weights, test correlations, ridge parameters and cross-validation splits are all saved in the specified ‘save_directory’. Finally, it computes the significance of the voxel correlations by running a blockwise permutation test from ‘significance_testing.py’.

Several of the utility functions imported in the script can be found in the ‘ridge_utils’ directory. ‘feature_spaces.py’contains implementations of each feature space—phoneme rate, articulators, word rate, English1000—and their corresponding downsampling functions. To add new feature spaces, one would need to implement a function that takes ‘story names’ as an argument and returns a dictionary of the downsampled stimulus features per story. Additionally, one would need to update the ‘_FEATURE_CONFIG’ variable. The ‘significance_testing.py’ script implements a parallelized block-wise permutation test which takes in a vector of predicted and true voxel responses, the number of permutations to test with and the size of each permutation block. It returns the associated p-value of each voxel. This script also contains a function to control for the false discovery rate using the Benjamini-Hochberg procedure.^26,44^

## Technical Validation (Unlimited Length)

One important metric for assessing BOLD data quality is head motion. Here we measured motion as framewise displacement^45^, which combines both rotation and translation into a single metric. **Figures 2A** and **2B** show framewise displacement for each subject and each story. **Figure 2A** shows the mean and range of average framewise displacement across all stories. This functions as a general metric for how still participants tend to be. Sub-UTS02 and sub-UTS03 have the lowest average displacement across stories while participants *Sub-UTS06-08* have the highest average displacement. **Figure 2B** shows the average displacement for each individual story in each participant. This shows that some of the participants with higher movement improved as the sessions went on and thus their high displacement values are due to outliers, as in *sub-UTS06* and *sub-UTS08*. The participant with the most consistent low framewise displacement is *sub-UTS02*. Out of a total 247 story scans across all subjects, the highest mean framewise displacement is only 0.33 mm.

**Figure 2.**
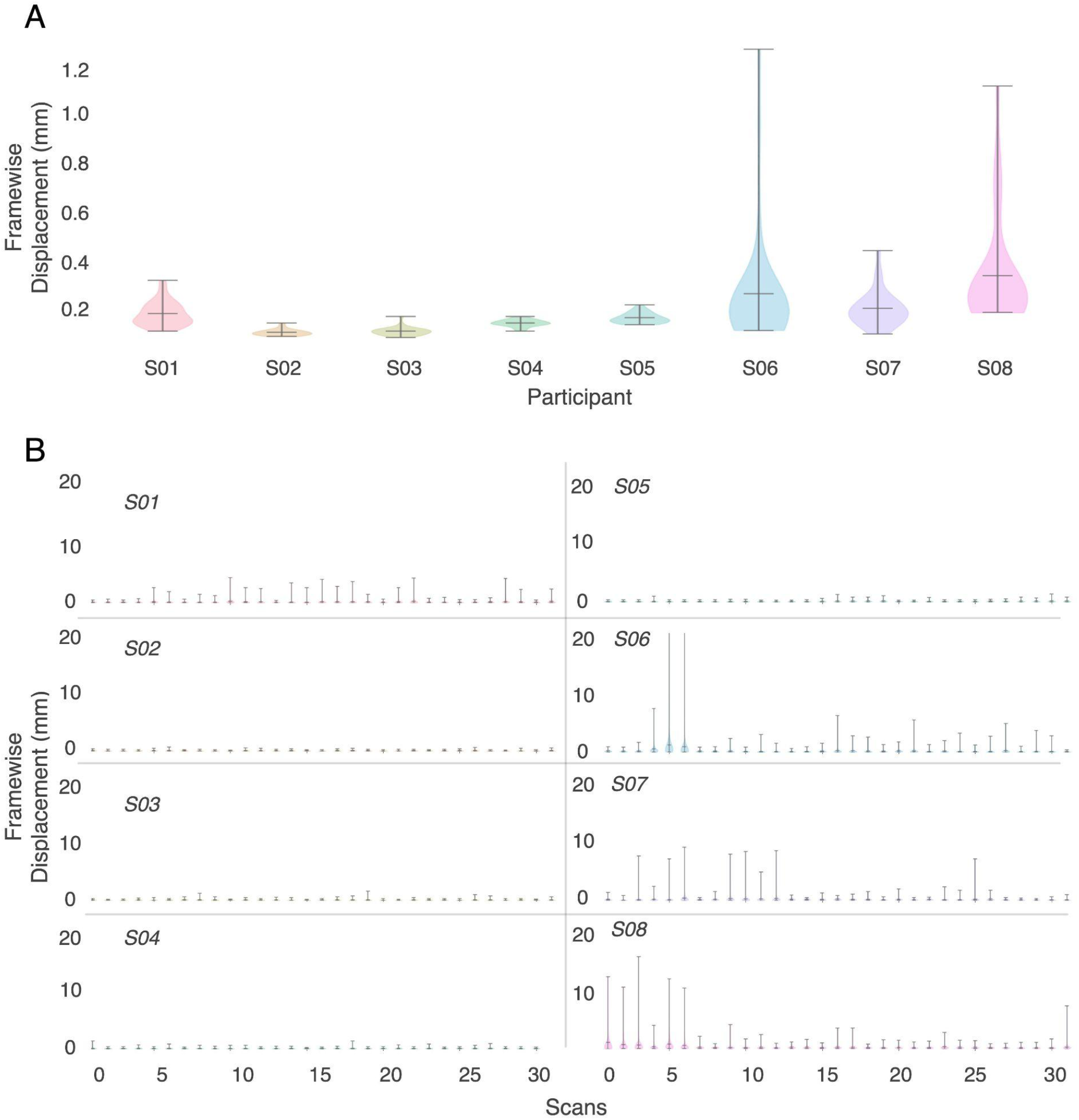
Head Motion across participants. Framewise displacement is measured as the mean shift needed to align each frame of data to the starting reference frame^45,46^. (A) Mean framewise displacement for each subject shows very low motion in participants S02-S05. Participants S06-S08 show the highest framewise displacement as they moved the most during data collection. (B) Mean framewise displacement is also assessed at the scale of each individual story for each subject. This similarly shows the lowest displacement for participants S02-S05 and the highest for participants S06-S08. However, these high movement participants had less motion over the course of data collection with later sessions having less movement.

Another important metric of fMRI data quality is functional repeatability, or how similar responses in the same voxel are to the same stimulus. While most of the stimuli were unique, the test story was played once in each of the five story sessions. This was done to enable us to compute the noise ceiling of the data and to increase the SNR of the held out dataset. It also enables us to measure how reliable the BOLD signal is in each voxel. Here, repeatability is calculated in each voxel as the mean pairwise correlation across the five validation story timecourses for that voxel, where a higher value means more reliable data. **Figure 3A** shows the repeatability for each subject as a step histogram across all voxels. The participants with the highest mean repeatability are sub-UTS03, sub-UTS02, and sub-UTS01.

**Figure 3.**
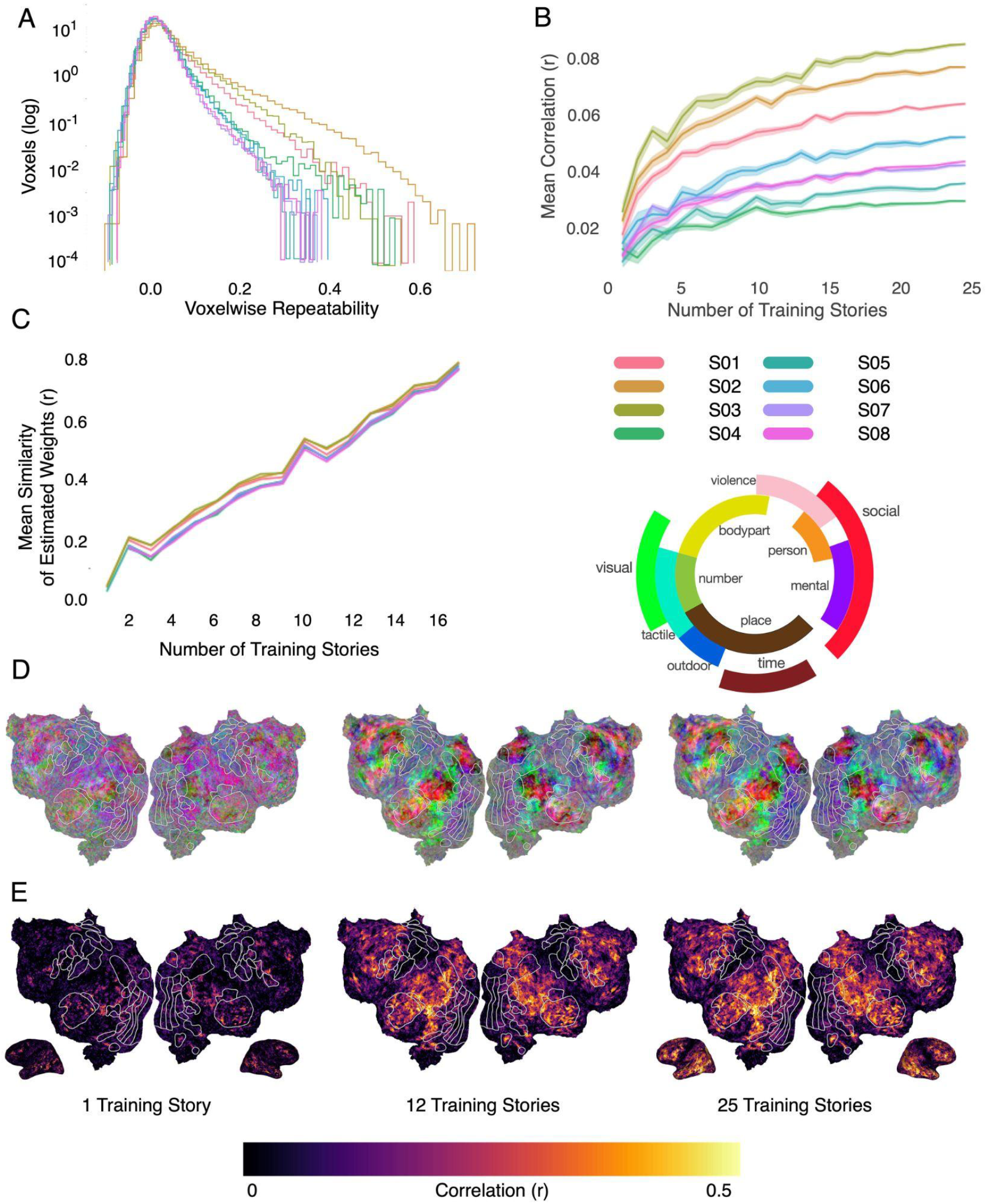
Model performance variance across subjects. (A) Voxelwise repeatability across five repeats of *task-wheretheressmoke* was calculated for each subject. Repeatability was calculated as the mean pairwise correlation across each repeat for each participant. Participants S01-S03 had the highest repeatability. (B) Word embedding-based encoding models^10^ were fit with increasing numbers of stories in the training set. As the dataset grows, so does model performance. Here we show the mean voxelwise model performance (r) for each subject. Participants S01-S03 had the highest model performance. (C) Using many stimuli for model training makes encoding weights stable, or invariant to the exact stimuli that were used. Here we measured weight stability by training encoding models using different stimulus subsets of varying sizes, and then computing the average pairwise correlation between the learned weights. As the training set grows, the estimated model weights for each voxel become more similar across different training subsets. (D) Encoding model weights were projected into a lower dimensional space^10^ to visualize the semantic map for one example participant (S02). As the training set grows, the semantic map appears to converge. (E) Encoding model performance—shown here projected onto a cortical flatmap for one participant—increases with the number of training stories. These increases are particularly evident in widespread temporal, parietal, and prefrontal cortex.

Lastly, one important metric used to assess data quality^12,13,43^ is encoding model prediction performance (r). In **Figure 3B**, we show the performance of semantic encoding models (mean(r)) as a function of the number of stories used for model training. For each number of training stories, we fit 15 models in which training data were sampled from the full set without replacement. Each point represents the mean prediction performance across all voxels and the cloud around the point represents the standard error across the 15 runs. All participants appear to reach a plateau where increasing the amount of training data does not dramatically improve performance. Note that this metric includes the majority of cortical voxels that are not semantically selective, biasing the result downwards. By this metric, the participants with the best data quality are sub-UTS03, sub-UTS02, and sub-UTS01. These subjects are also the subjects with the lowest motion and highest repeatability. **Figure 3E** shows the voxelwise prediction performance (r) for one subject plotted on the flatmaps for an increasing number of stories. This again shows that as the amount of data increases, more voxels are predicted and predicted better.

This improvement in model performance means that the model weights from the fit encoding models become more stable and less affected by attention and noise (**Figure 3C**). To demonstrate the importance of large datasets within an individual for interpreting model weights, we fit models using different subsets of stories and then calculated the mean pairwise correlation across estimated model weights. As the number of stories in the subsets increases, the estimated model weights become more similar regardless of the individual stories used in training. This makes interpretation of the model weights more robust. **Figure 3D** shows the model weights for each of these models projected into a three-dimensional semantic space that was previously constructed from a group of subjects using principal components analysis.^10^ This lower-dimensional space is used purely for visualization purposes. Here projections on the first, second, and third principal components are mapped into the red, green, and blue color channels, respectively, for each voxel and then projected onto the cortical surface. The color wheel shows approximately which semantic category each color on the maps represents. The separability and intensity of the weight increases and becomes clearer as dataset size grows. Increasing the size of datasets within individuals thus increases reliability and interpretability of encoding models, and is vital to increasing the reliability of results in the field.

## Usage Notes

Anatomical and localizer scans (ses-1) for sub-UTS02 were collected prior to this current dataset, at a different location, and with a different scan protocol than all other data in this project. Consequently, the localizer data is not included here. However, the hand-defined regions of interest (ROIs) derived from the localizer data can be found for this participant in the pycortex-db in their overlays file. This participant also has additional ROIs from other localizers including retinotopy.

Participant sub-UTS04 has one missing story scan, task-life.

Participant sub-UTS05 was presented with auditory cues for the motor localizer stimuli instead of visual cues. Additionally, this localizer lacked the saccade cue. This difference was due to the participant’s visual acuity being too low to successfully read the cue and MRI-safe glasses were not compatible with the headcase being used.

## Code Availability

All code used for encoding model fitting is publicly available and can be found at https://github.com/HuthLab/deep-fMRI-dataset.

## Notes

This work was supported by the Whitehall Foundation, Alfred P. Sloan Foundation, Burroughs-Wellcome Fund, and the Texas Advanced Computing Center (TACC). The Authors Declare no conflict of interest.

https://openneuro.org/datasets/ds003020

https://github.com/HuthLab/deep-fMRI-dataset

### Competing Interest Statement

The authors have declared no competing interest.

